# Dynamics in vertical transmission of viruses in naturally selected and traditionally managed honey bee colonies across Europe

**DOI:** 10.1101/2022.03.25.485775

**Authors:** David Claeys Bouuaert, Lina De Smet, Marleen Brunain, Bjørn Dahle, Tjeerd Blacquière, Anne Dalmon, Daniel Dezmirean, Dylan Elen, Janja Filipi, Alexandru Giurgiu, Aleš Gregorc, John Kefuss, Barbara Locke, Joachim R. de Miranda, Melissa Oddie, Delphine Panziera, Melanie Parejo, M. Alice Pinto, Dirk C. de Graaf

**Affiliations:** Department of Biochemistry and Microbiology, Ghent University, Ghent, Belgium; Faculty of Environmental Sciences and Natural Resource Management, Norwegian University of Life Sciences, Ås, Norway; Bees@wur, Wageningen University & Research, Wageningen, The Netherlands; Abeilles et Environnement, INRAE, Avignon, France; Department of Apiculture and Sericulture, University of Agricultural Sciences and Veterinary Medicine, Cluj-Napoca, Romania; Department of Molecular Ecology & Evolution, School of Natural Sciences, Bangor University, Bangor, United Kingdom; Taskforce Research, ZwarteBij.org VZW, Gavere, Belgium; Department of Ecology, Agronomy and Aquaculture, University of Zadar, Zadar, Croatia; Faculty of Agriculture and Life Sciences, University of Maribor, Pivola, Slovenia; Le Rucher D’Oc, Toulouse, France; Department of Ecology, Swedish University of Agricultural Sciences, Uppsala, Sweden; Applied Genomics and Bioinformatics, University of the Basque Country, Leioa, Spain; Centro de Investigação de Montanha, Instituto Politécnico de Bragança, Campus de Santa Apolónia, Bragança, Portugal; Norwegian Beekeepers Association, Kløfta, Norway

## Abstract

The ‘suppressed *in-ovo* virus infection’ trait (SOV) was the first trait applied in honey bee breeding programs aimed to increase resilience to virus infections, a major threat for colony survival. By screening drone eggs for viruses, the SOV trait scores the antiviral resistance of queens and its implications for vertical transmission. In this study, queens from both naturally surviving and traditionally managed colonies from across Europe were screened using a two-fold improved SOV phenotyping protocol. First, a gel-based RT-PCR was replaced by a RT-qPCR. This not only allowed quantification of the infection load but also increased the test sensitivity. Second, a genotype specific primer set was replaced by a primer set that covered all known deformed wing virus (DWV) genotypes, which resulted in higher virus loads and fewer false negative results. It was demonstrated that incidences of vertical transmission of DWV were more frequent in naturally surviving populations than in traditionally managed colonies, although the virus load in the eggs remained the same. Dynamics in vertical transmission were further emphasized when comparing virus infections with queen age. Interestingly, older queens showed significantly lower infection loads of DWV in both traditionally managed and naturally surviving colonies, as well as reduced DWV infection frequencies in traditionally managed colonies when compared with younger queens. Seasonal variation in vertical transmission was found with lower infection frequencies in spring compared to summer for DWV and black queen cell virus. Together, these patterns in vertical transmission suggest an adaptive antiviral response of queens aimed at reducing vertical transmission over time.

## Introduction

Disease pressure is an inherent driver in the evolution of eusociality [1,2] and social task division [3] and thus forms an important component in the evolution of Western honey bees (*Apis mellifera*). Before the arrival of the Varroa mite (*Varroa destructor*), virus infections were mostly benign and rarely associated with colony losses [4]. The arrival of the Varroa mite considerably changed the virus landscape by introducing a new transmission pathway and thereby influencing virus virulence and evolution [5–13]. Through varroa-mediated transmission, virus diseases have become one of the most important proximate causes of colony mortality and honey bee decline [14–21].

Of the 72 virus species that have been identified in honey bees [22], the most commonly occuring belong to *Iflaviridae* and *Dicistroviridae* [23,24], particularly sacbrood virus (SBV), black queen cell virus (BQCV), acute bee paralysis virus (ABPV) and deformed wing virus (DWV), with both ABPV and DWV consisting of a complex of closely related, co-circulating master variants capable of forming viable recombinants [22,25,26]. The DWV virus complex is best described as a group of functionally and genetically compatible minor and major variants and their recombinants based on four master strains [27], of which DWV-A and DWV-B are currently the most common [28–33]. Dynamics in the presence and abundance of honey bee viruses show strong seasonal and geographic variation [34–36]. This variation is driven by local adaptation of both virus, host and vector species as well as by the specific characteristics of each virus [37–41]. Together they form a geographic mosaic of coevolution [42].

In the first years after managed colonies are left untreated against the Varroa mite, colony mortality increases considerably [43,44]. This results in strong selective pressure forcing bees, mites and the viruses to adapt to each other. Most naturally surviving populations consist of unmanaged or feral colonies [41]. In managed colonies, two approaches have been described to transition from treated colonies to naturally surviving colonies. The first consists of leaving a large number of colonies unmanaged with respect to swarming, re-queening and varroa control [40] and allowing natural selection to take place. This is described as the ‘Bond’ test: ‘Live and let die’ [40]. A second approach, named ‘Darwinian black box’ builds further on this by adding closed population mating and selection for strong spring development [44]. One of the best studied naturally surviving populations with regard to virus-host coevolution is an isolated, closed honeybee population located at the tip of Näsudden peninsula in the south of Gotland, a Swedish island in the Baltic sea. After implementing the Bond test, these honey bee colonies evolved an increased tolerance for DWV infections [39,45]. In addition, BQCV and SBV infections were less abundant in autumn and early spring, possibly due to the reduced colony size of the Gotland colonies in these seasons [46].

In general, coevolution tends to evolve towards a maximal investment in defense strategies based on increased tolerance or resistance in the host and an intermediate virulence in the virus [47]. In the case of the Gotland population, this maximal investment in defense strategies resulted in increased virus tolerance [39] and adapted colony demographics [46]. By definition, tolerance strives to minimize the cost of the infection, resistance aims to prevent infection. In contrast to tolerance, resistance will reduce the prevalence of the parasite in a population [48–50] and impose selection on the parasite to overcome the host defense, which can eventually lead to antagonistic coevolution [51]. In a honey bee colony, resistant individuals will lead to a resistant colony. If only parts of the individuals are resistant, the colony as a whole may become tolerant to the parasite, as has been shown in the Varroa tolerant colonies in Gotland [52]. In contrast, a colony composed of tolerant individuals may become resistant to the parasite. This was shown in a Nosema-tolerant breeding line in Denmark [53]. Individual bees developed high infection loads but without metabolic energy costs. This was suspected to result in reduced transmission between individuals and eventually led to the clearance of the infection [53]. In a honey bee colony, tolerance and resistance can thus act at both the individual and the colony level. Important to take into account is that both strategies have possible associated costs and can often coexist [54–56].

Each response to a parasite influences transmission dynamics within and between honey bee colonies. As honey bees live closely together with thousands of individuals, the transmission of viruses through trophallaxis, feeding or body contact occurs frequently. This form of transmission between individuals of the same generation is defined as horizontal transmission. On the other hand, transmission between generations by either eggs or semen is defined as vertical transmission [26]. Virus infections of queens, or their eggs, have been shown to interfere with normal egg development, to elicit a stress response in eggs [57], and to cause important health risks for the queen herself [58–61]. The importance of the honey bee queen in the viral dynamics of the colony was highlighted with the discovery of the ‘suppressed *in-ovo* virus infection’ trait (SOV). Colonies headed by a queen laying virus-free eggs showed fewer and less severe DWV infections in almost all developmental stages of both drones and workers. In addition, this potential to suppress viral infections is heritable [62] and alters the tissue specificity of DWV [63].

The aim of this study is to screen for the presence of the SOV trait in naturally surviving (NSC) and traditionally managed colonies (TMC) across Europe, along with the analysis of the effect that queen age and the time during the bee season have on the SOV trait. To this end, two improvements were made to the phenotyping protocol for the SOV trait [62]. First, the detection of viral pathogens by RT-qPCR instead of gel-based RT-PCR allows for a quantification of the viral load of the eggs and lowers the detection threshold. Second, as the SOV trait is associated with increased virus resistance across DWV genotypes [63], a shift was made from screening for DWV-A only to a generic screening for the DWV complex. Overall, this research improves our understanding of how patterns of vertical transmission of viruses differ across Europe and in different evolutionary settings.

## Methods

### SOV phenotyping

Samples collected as part of the Flemish bee-breeding program in 2020 were used to compare the transition from a gel-based RT-PCR to RT-qPCR approach and to compare the quantification of DWV infections using primers specific to DWV-A or DWV-B and a generic DWV primer (DWV-Fam). All samples were collected following the phenotyping protocol for the SOV trait, as described by de Graaf, et al. (2020), and were screened by RT-qPCR for DWV-A, DWV-B, DWV-Fam, SBV, ABPV and BQCV. For comparing the performance of the gel-based RT-PCR with the RT-qPCR, positive samples covering a 10^1^-10^8^ / 10 eggs range were selected and analyzed by gel-based RT-PCR for SBV, ABPV, DWV-A and BQCV.

### Egg sample collection across Europe

Egg samples were collected in 9 countries across Europe during spring and summer 2020. Each country sampled 10 TMC (colonies managed following local standard practices including treatment against the Varroa mite) and, if present, 10 NSC (colonies from populations that survive without treatment against the Varroa mite). Colonies were managed following local standard practices and queens descended from locally adapted or native stock. Treatment of the TMC colonies against the Varroa mite was performed with registered products in each country. Management for the NSC was conducted according to the ‘Darwinian Black Box’ selection method [44] or the ‘Bond’ test [40]. From each queen, a pooled sample of 10 drone eggs was collected following the phenotyping protocol of the SOV trait, as described by de Graaf et al (2020). If drone eggs were not present and attempts to induce drone laying did not succeed, worker eggs were collected instead. All samples were immediately stored at −20 °C and kept in a cold chain during transport to Belgium, where they were analyzed for DWV-Fam, BQCV, SBV and ABPV by RT-qPCR. For each sampled colony, information was gathered on the sampling season, subspecies, queen age, beekeeping method (for both TMC and NSC) and colony health status at the time of sampling. This information was used to explain possible outliers and to look for correlations between multiple factors. Additional samples were collected if the apiary was composed of more than 10 colonies and if drone eggs were present during sampling. Samples from Slovenia were collected in the scope of a different project, hence the larger sample size.

### Gel-based RT-PCR

All samples were first homogenized in the presence of zirconium beads in 0.5 ml QIAzol lysis reagent (Qiagen). RNA was extracted using the RNeasy Lipid tissue mini kit (Qiagen) according to the manufacturer’s instructions, including a DNAse step, and finally eluted in 30 μl elution buffer. The concentration of the total RNA was measured with Nanodrop (Isogen). Using random hexamer primers, 200 ng RNA was retro-transcribed with the RevertAid H Minus First Strand cDNA Synthesis Kit (Thermo Scientific). Honey bee β-actin was used to control RNA integrity. All gel-based RT-PCR reaction mixtures contained: 2 μM of each primer (see Supplementary S1); 1 mM MgCl_2_; 0.2 mM dNTPs each; 1.2 U HotStarTaq Plus DNA polymerase (Qiagen) and 2 μl cDNA product. Gel-based RT-PCR assays were performed using the following cycling conditions: 95°C - 5 min; 94°C – 30 sec, 55°C - 30 sec, 72°C - 1 min, 35 cycles; final elongation 72°C - 10 min, hold 4°C. Gel-based RT-PCR amplicons were analyzed by electrophoresis using 1.5% agarose gels stained with ethidium bromide and visualized under UV light. Positive and negative controls were included in each run.

### RT-qPCR

The virus load RT-qPCR determination was performed using Platinum™ SYBR™ Green qPCR SuperMix-UDG (Thermo Scientific). Each reaction consisted of 0.4 μM of each primer (sequences provided in Supplementary S1), 11.45 μl RNase-free water, 12.5 μl SYBR Green and 1 μl of cDNA template. All samples were run in duplicate in a three-step RT-qPCR. Thermal cycling conditions started with an initial activation stage at 95 °C for 2 min followed by 35 cycles of a denaturation stage at 95°C for 15 sec, annealing stage at 58°C for 20 sec and extension stage at 72°C for 30 sec. This procedure was followed by a melt-curve analysis to confirm the specificity of the product (55–95 °C with an increment of 0.5 °C sec−1). Each plate included a no template and a positive control. A standard curve obtained through an 8-fold 5x serial dilution of a known amount of viral plasmid loads (range of 10^4^ – 10^10^ copies / μl) was used for absolute quantification. All data were analyzed using CFX Manager™ 3.1 software (Bio-Rad). Baseline correction and threshold setting were performed using the automatic calculation offered by the same software. Maximum accepted quantification cycle (Ct) difference between replicates was set to two Ct. The successful amplification of the β-actin internal reference gene was used to confirm RNA integrity throughout the entire procedure [64].

### Statistics

Viral loads for each sample were log10-transformed to improve data visualization. Detection thresholds for all pathogens were set at 30 Ct (corresponds to 10^3^ copies for DWV, BQCV, SBV and APBV). Below this threshold, samples cannot be reliably quantified by RT-qPCR [65]. RStudio version 3.6.1 was used for data analysis and visualization. Analyses of the differences in the number of infections between groups were conducted by Chi-squared tests. For comparison between the infection loads T-tests were used. All tests were checked for and complied with the required assumptions.

## Results

### SOV phenotyping method

The virus detection threshold on gel-based RT-PCR (based on a RT-PCR of samples positive on RT-qPCR along a 10^1^-10^8^ copies / reaction range of starting template) is 10^2^ for DWV-A, 10^7^ for BQCV, 10^6^ for ABPV and 10^8^ for SBV (see Supplementary S2). A total of 187 samples were screened for DWV-A, DWV-B and DWV-Fam (generic DWV primer). Of these, 153 (82%) showed amplification when screening with DWV-Fam. One sample amplified with just the DWV-A assay and two samples with just the DWV-B assay. Of the 153 samples that amplified with the DWV-Fam assay, 98 (64%) also amplified with the DWV-B assay, two (1%) with the DWV-A assay and six (4%) with both the DWV-A and DWV-B assays. The remaining 47 samples (25%) only amplified with the DWV-Fam assay and not with either the DWV-A or DWV-B assay. The median infection load for DWV-Fam was 5.8 Log10 virus copy number / egg and was on average 1.69 Log10 higher compared to the sum of DWV-A and DWV-B.

### European SOV phenotyping

Table 1 shows the number of collected samples, the number of infections for each virus, and the mean infection load for each country and for each selection strategy (TMC or NSC). Of the 213 egg samples screened, most infections were with DWV (51%), followed by BQCV (25%). Only three samples were infected with SBV and two samples were infected with ABPV. Differences in infection frequencies varied considerably between and within countries. Supplementary Table S3 provides an overview of the sampled populations and countries, the number of worker egg samples, the colony health status at the time of sampling, and lists the location of the sampled populations. Worker egg samples from Norway and Sweden had lower infection frequencies (11/34) than drone egg samples from other locations (61/107). Slovenian samples were collected in the scope of a different project, hence the larger number of samples. To avoid uneven distribution of the sample size across groups, Slovenian samples were not included during further analyses. Differences between subspecies could not be analyzed due to the high variability between countries and hybridization between subspecies. Due to bad weather conditions or the inaccessibility of some locations some countries (Netherlands, Romania and Sweden) were not able to sample the requested number of colonies.

**Table 1:** Overview of SOV phenotyping results for both naturally surviving (NSC) and traditionally managed (TMC) colonies in each participating country. SOV+ indicates the number of egg samples free of virus infections.

| Country | NSC /<br>TMC | Nr. of<br>samples | SOV+ | Nr. of samples positive for: |  |  |  | Mean infection load (Log10 / egg) |  |  |  |
| --- | --- | --- | --- | --- | --- | --- | --- | --- | --- | --- | --- |
|  |  |  |  | DWV | BQCV | SBV | ABPV | DWV | BQCV | SBV | ABPV |
| Belgium | NSC | 10 | 1 | 9 (90%) | 0 | 0 | 0 | 4.3 |  |  |  |
|  | TMC | 11 | 4 | 7 (64%) | 2 (18%) | 0 | 0 | 6.1 | 4.2 |  |  |
| Croatia | TMC | 10 | 3 | 4 (40%) | 3 (30%) | 1 (10%) | 0 | 4.7 | 5.3 | 3.3 |  |
| France | NSC | 13 | 2 | 7 (58%) | 11 (85%) | 0 | 0 | 5.6 | 5.5 |  |  |
|  | TMC | 10 | 2 | 1 (10%) | 8 (80%) | 0 | 0 | 5.8 | 6.4 |  |  |
| Netherlands | NSC | 10 | 2 | 8 (80%) | 3 (30%) | 0 | 0 | 6.2 | 5.4 |  |  |
|  | TMC | 6 | 1 | 5 (83%) | 3 (50%) | 0 | 0 | 5.4 | 4.9 |  |  |
| Norway | NSC | 10 | 1 | 9 (90%) | 1 (10%) | 0 | 0 | 5.1 | 4.9 |  |  |
|  | TMC | 10 | 5 | 4 (40%) | 3 (30%) | 0 | 0 | 4.3 | 4.7 |  |  |
| Portugal | TMC | 10 | 1 | 8 (80%) | 1 (10%) | 0 | 0 | 5.1 | 3.3 |  |  |
| Romania | NSC | 4 | 1 | 0 | 3 (75%) | 0 | 0 |  | 4.5 |  |  |
|  | TMC | 9 | 4 | 2 (22%) | 4 (44%) | 0 | 0 | 4.0 | 4.0 |  |  |
| Slovenia | TMC | 72 | 27 | 38 (53%) | 11 (15%) | 2 (2%) | 2 (2%) | 5.3 | 5.2 | 3.2 | 3.8 |
| Spain | TMC | 10 | 4 | 5 (50%) | 1 (10%) | 0 | 0 | 5.5 | 4.8 |  |  |
| Sweden | NSC | 6 | 5 | 1 (16%) | 0 | 0 | 0 | 5.0 |  |  |  |
|  | TMC | 12 | 11 | 1 (8%) | 0 | 0 | 0 | 4.3 |  |  |  |

Figure 1 shows the distribution of the number of virus infections in TMC for each participating country. Queens laying virus-negative eggs score positively on the SOV trait (SOV+) and are represented by the green fraction (labelled as SOV positive in figure). Sweden had the highest percentage of SOV+ queens notwithstanding the presence of clinical symptoms of DWV, chalkbrood and possibly EFB visible during sampling. With the exception of colonies in the Netherlands, most countries showed limited numbers of multiple virus infections in the same sample (Figure 1).

**Figure 1:**
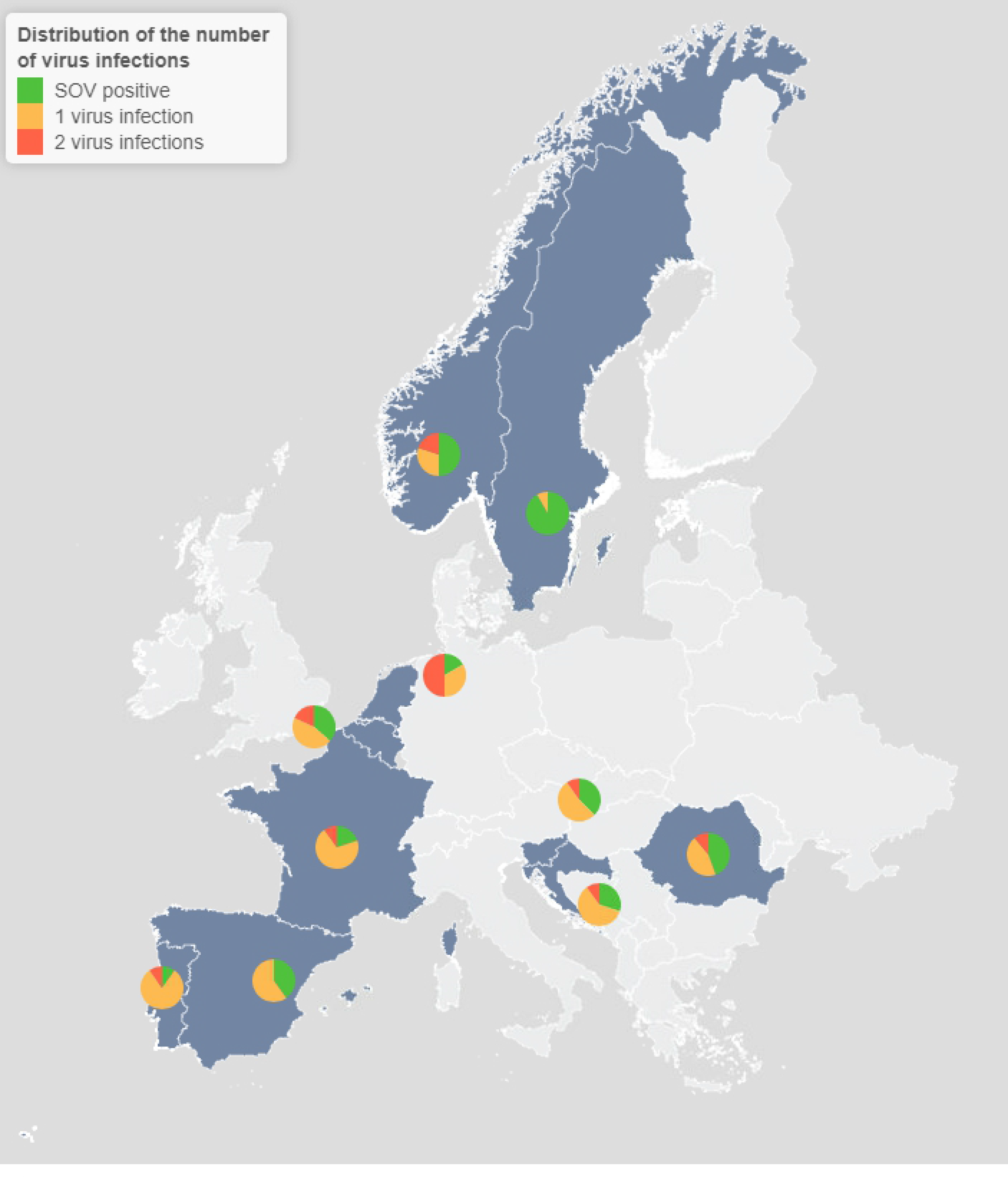
Distribution of the number of virus infections per sample in traditionally managed colonies for each country. SOV positive indicates samples free of virus infections.

Figure 2 shows the percentage of virus infections occurring in different parts of the season. Samples collected in spring had significantly higher infection frequencies than samples collected in summer for both DWV (X^2^(1, N=141) = 9.4, p < 0.05) and BQCV (X^2^(1, N=141) = 12.3, p < 0.05) but not for SBV.

**Figure 2:**
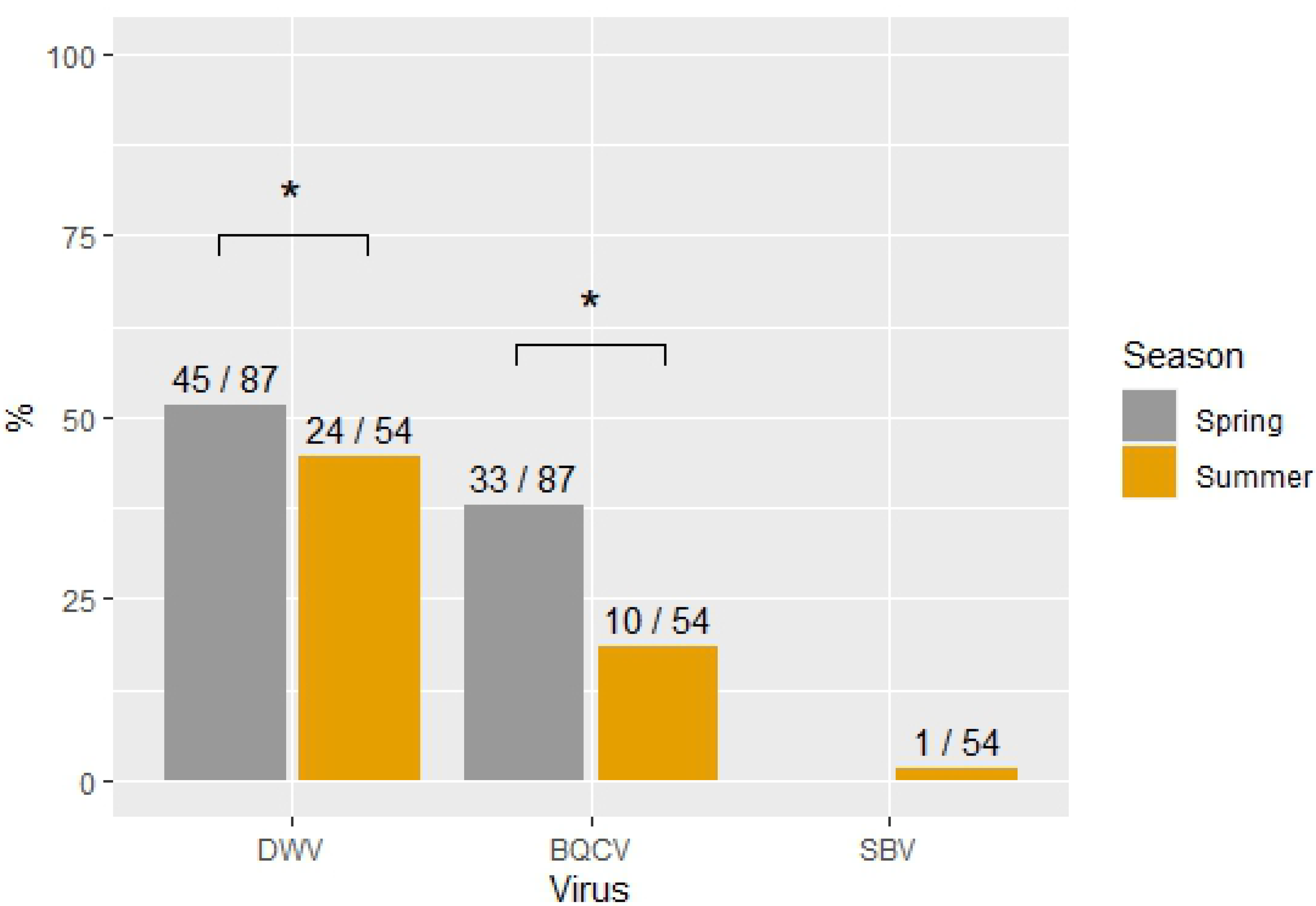
Percentage of virus infections in each sampling season. Infections were significantly more frequently in samples collected in spring for DWV and BQCV as compared to samples collected in summer. Significant differences are indicated with *.

### Natural survivors vs traditionally managed colonies

Figure 3 shows the percentage of virus infections (A) and the infection loads (B), for both NSC and TMC, and for each virus. Infection frequencies were significantly higher in NSC compared to TMC for DWV (X^2^(1, N=111)= 8.6, p < 0.05) but not for BQCV (X^2^(1, N=111) = 0.1, p = 0.75). No significant differences were found between the infection loads of TMC and NSC for DWV (t(68) = −0.6, p = 0.52; TMC = 5.2 ± 0.4, NSC = 4.9 ± 1.6) and BQCV (t(33) = 1.6, p = 0.11; TMC = 4.4 ± 0.9, NSC = 4.8 ± 0.1). On average, most infections hovered around 5 Log10 for both DWV and BQCV.

**Figure 3:**
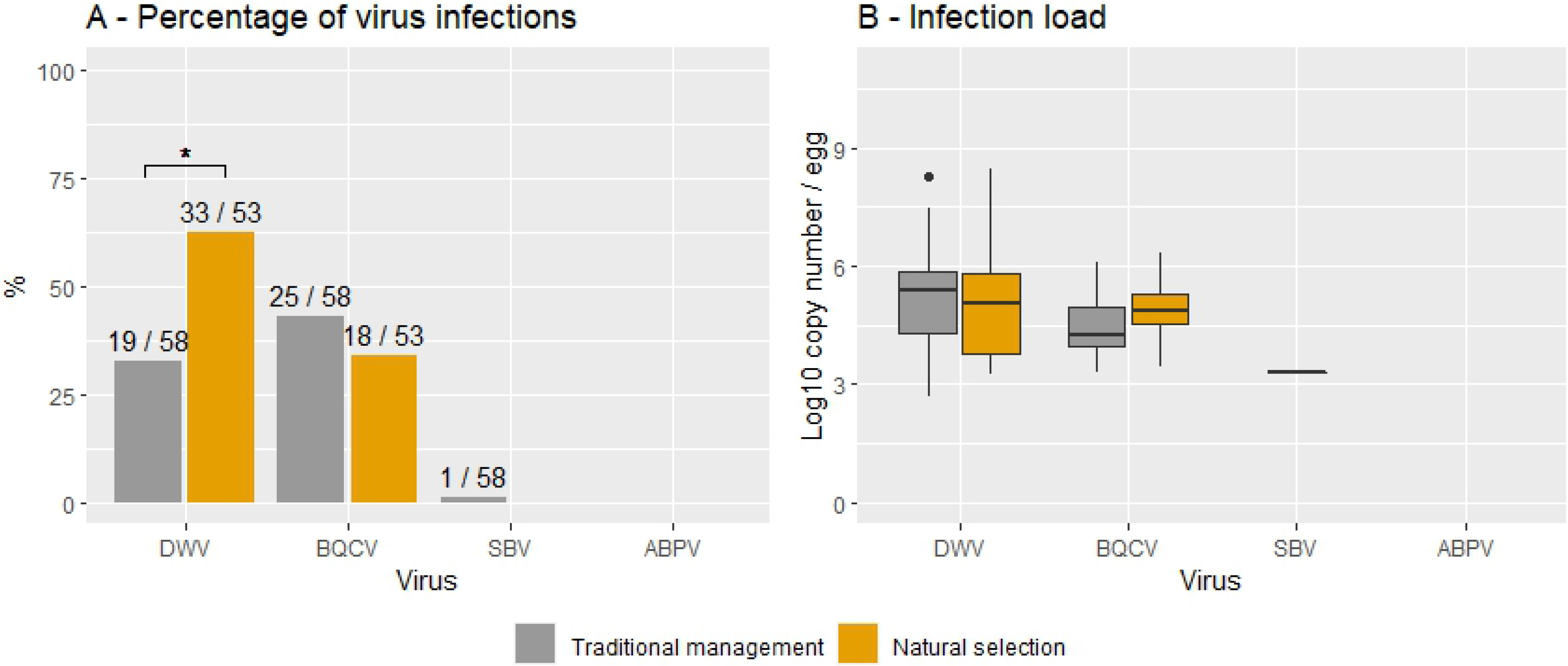
Percentage of viral infections (A) and infection loads (B) for naturally selected and traditionally managed colonies. *Data is provided for each virus.* Significant differences are indicated with *.

### Queen age

Figure 4A shows the frequency of infection with DWV or BQCV for each queen age and for both TMC and NSC groups. For DWV, a significant decrease in the percentage of infected samples was found in TMC between queens aged 0 and 1 year (X^2^(1, N=59) = 3.9, p < 0.05). This trend continued with a lower infection frequency in queens aged 2 years (1/10) compared to queens aged 1 year (14/42), albeit not being significant. No differences in infection frequencies were found between queen ages in NSC. Figure 4B shows the infection loads of DWV and BQCV for each queen age for both TMC and NSC groups. As previously shown, the infection load did not differ between the two groups. Comparing infection loads between queen ages showed significantly higher infection loads in queens aged 0 years (M = 5.7, SD = 0.2) compared to queens aged 1 year for DWV (M = 4.8, SD = 1.6; t(29) = 2.6, p < 0.05) for both TMC and NSC. Albeit not being significant, the mean DWV infection load for queens aged 2 years (M = 5.3) was lower than the mean of queens aged 1 year (M = 6.2). No significant differences in infection frequencies nor infection load were found for BQCV. The infection load did however show a similar general trend, with decreasing mean infection loads with age (5.0 in queens aged 0 years, 4.6 in queens aged 1 year and 4.4 in queens aged 2 years). Interestingly, the spread in infection loads of queens aged 0 largely lacks infection loads lower than 10^7^ for DWV and BQCV. Seasonal differences in sample collection did not influence the infection frequency for DWV in queens aged 0 years (X^2^(1, N=25) = 1.69, p = 0.16) and older queens (X^2^(1, N=97) = 2.2, p-value = 0.14). Queen age was unknown for 19 out of 141 samples (13%).

**Figure 4:**
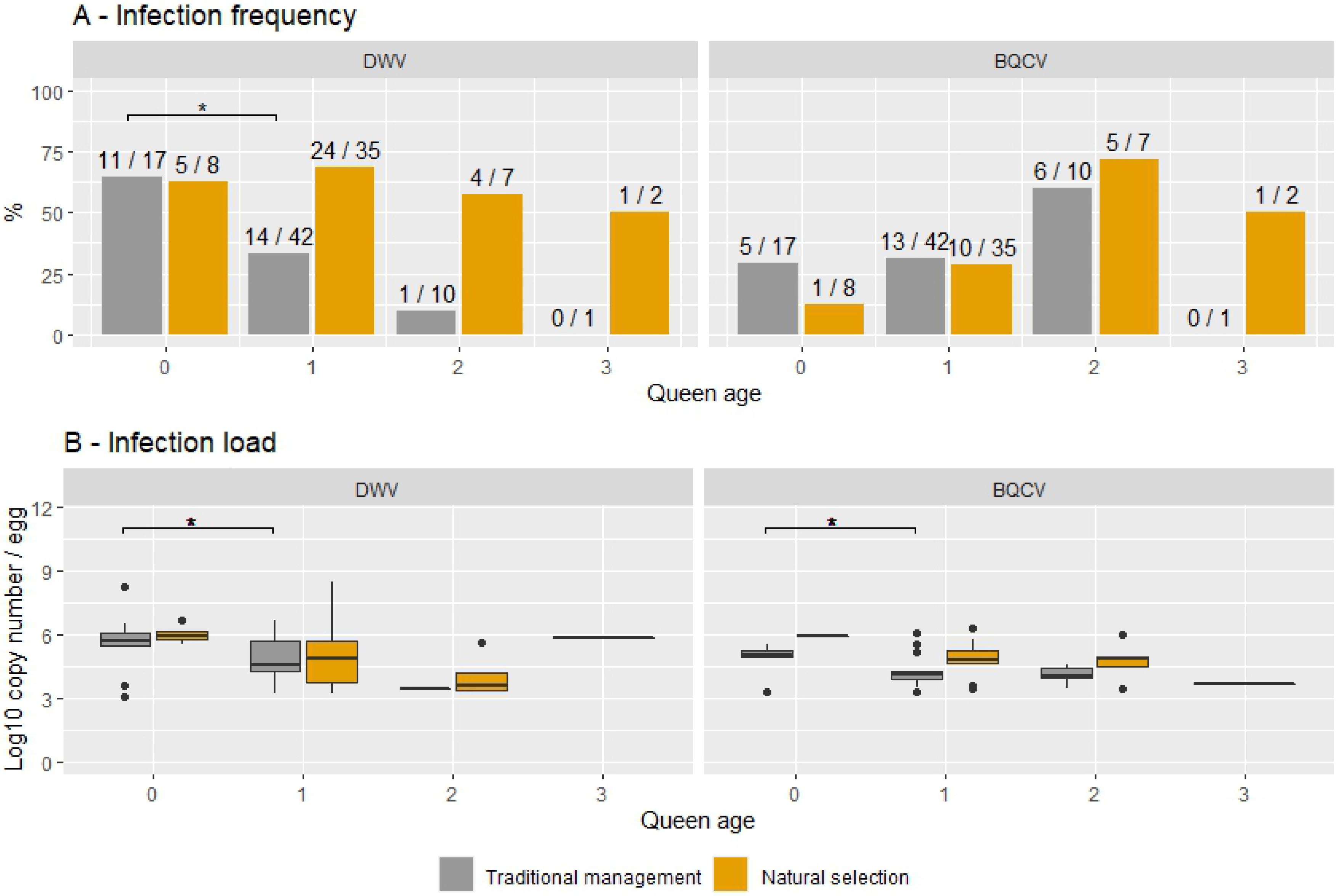
Overview of the infection frequency (A) and the infection load (B) for each queen age from both naturally selected and traditionally managed colonies. Significant differences are indicated with *.

## Discussion

Phenotyping for the SOV trait by RT-qPCR showed, as expected, a higher detection sensitivity as compared to gel-based RT-PCR. The detection threshold for RT-qPCR was around 10^3^ (30Ct) for all viruses, while gel-based RT-PCR showed significantly higher detection thresholds for SBV, BQCV and ABPV (10^8^ for SBV, 10^7^ for BQCV and 10^6^ for ABPV). This implies that phenotyping by gel-based RT-PCR underestimated the number of virus infections for these viruses. There was no difference between both detection thresholds for DWV. It should be noted that samples negative on RT-qPCR can still be infected below the detection threshold and that positive samples might be infected with viruses in a dormant state. An important advantage of RT-qPCR is that breeding programs can manually set threshold values for SOV phenotyping based on the breeding goal and the virulence of the virus. Each breeding program can thus determine the degree of positive or negative selection desired. Infection loads in eggs are linked with the infection status of the queen [66,67] and have been shown to reduce virus infections in the colony as a whole [62]. Nevertheless, the impact of different virus infection loads in eggs on subsequent developmental stages is currently unknown. Further research, where eggs with different virus infections are reared *in-vitro,* could improve our understanding of the impact that vertical transmission has on antiviral responses and honey bee health.

The comparison between individual DWV genotypes and the generic DWV shows a large underestimation of DWV infections when screening for either one of the genotypes or the sum of DWV-A and DWV-B. This can be seen in terms of an underestimation of the number of infected samples (25% of the samples) and lower infection loads (on average 1.69 Log10 lower). In comparison, a previous study in the UK found 40% higher DWV titers when screening with a universal DWV-complex assay than by pooling the results of screening with specific DWV-A and DWV-B assays [68]. Possible explanations can be the presence of different genotypes, DWV-C or DWV-D, neither of which has so far been detected in Belgium [27,28], or mutations in the primer region that hamper correct primer hybridization [69]. A study on the genetic diversity within a DWV population in a colony showed that 82% of the genome had >1 sequence variant present in the frequency of >1%, and 39% of the genome had >1 sequence variant present in a frequency of >10% [8]. In addition, shifts in the sequence space of the DWV-A quasispecies have been shown after injection in a honey bee pupae [70]. The rapid shifts in the DWV quasispecies are consistent with the punctuated evolution theory, whereby infection of a new host causes a selective sweep followed by diversification towards an increased genetic heterogeneity that has potentially adapted to the host specific antiviral defenses [29]. This implies that primer regions, although being in conserved regions, may evolve over time and reduce the primer amplification efficiency.

Queens laying SOV+ eggs (free of viruses) were found in all countries and in both TMC and NSC groups. Remarkable was the low number of infections in Sweden where both groups only had one sample infected with DWV, despite multiple studies recording high viral loads in the worker bees in this population throughout the years [36,39,45,46,71–73]. The presence of the SOV trait across Europe serves as a possible starting point for local breeding programs to perform selection within the variation in virus resistance present within honey bee populations [62,74]. In addition to the high prevalence of DWV, the second most common virus found in this study was BQCV. This virus is the most common cause of queen larval death [75,76] but has not been found to cause overt symptoms in queens despite the detection of high infection loads [77]. Viruses of the ABPV complex and SBV have been found in eggs [62,67,78,79] but were rarely detected in this study. The high virulence of BQCV, SBV and ABPV [4,61,80–83] compared to the low virulence of DWV [80] could explain why they are less likely to be transmitted vertically without causing queen supersedure or colony health issues [84].

Differences in infection patterns between countries can be caused by climatic conditions, the seasonality of honey bee viruses [35,69,85–91] or by the low number of samples per country. This is reflected in the significant differences in infection frequencies between the spring and summer sampling seasons in this study. Interestingly, infection frequencies with DWV were higher in spring, despite the generally higher infection frequencies in adult bees during summer and autumn [86,88,91]. These findings implicate that conclusions based on SOV phenotyping should always take the time of sampling into account when interpreting results. Other factors affecting virus abundance are nutritional quality and availability [92,93], the connectivity between colonies [94], colony demography [95–97], population heterogeneity [98–102], colony management [103], degree of local adaptation [104], the individual and colony-level immune responses [105] and other stress factors such as exposure to neonicotinoids [106]. In this study worker eggs did not have higher infection loads compared to drone eggs despite the possible occurrence of trans-spermal virus transmission [26].

By comparing naturally surviving with traditionally managed populations in the same local context, insights can be gained into which evolutionary adaptations are needed for honey bees to survive without treatment against the Varroa mite. Typical for naturally surviving populations is that they harbor higher mite numbers that serve as an important vector for viruses [40,107]. Honey bee colonies react to these high disease pressures with adaptations in their antiviral responses or by forms of social immunity [108]. This study shows that infection frequencies were significantly higher in NSC than in TMC for DWV but that infection loads did not differ between the two groups. Honey bee queens appear to avoid increased vertical transmission loads despite increased infection frequencies. Virus loads in worker bees were not studied here. Therefore, it remains uncertain if the higher infection frequencies are a result of increased virus circulation in the naturally surviving populations. The high variability between both groups in each country can be caused by previously mentioned factors affecting virus abundance, the time since colonies were left untreated [37,38], or the degree of genetic divergence between TMC and NSC within a country [62,74,102].

Honey bee queens accumulate viral infections and infection loads during queen rearing [109,110], during mating flights [58,60,111] or as they get older [60,64,79] despite increased immune responses [112]. In contrast to what was expected, both the infection frequency of DWV and the infection load in eggs of queens from TMC decreased with increasing queen age. This difference is not caused by the mortality of queens with high infection loads as the distribution of the egg infection loads does not overlap between queens aged 0 and 1 year. For BQCV a similar trend was found but did not show significant differences. Queens from NSC showed the same trends in infection loads between all queen ages but did not show differences in infection frequencies. In beekeeping practices, queens are often renewed yearly as young queens are associated with lower winter mortality [113,114]. This study suggests that older queens are able to adapt their antiviral responses and reduce the infection loads transmitted via their eggs. If so, frequent renewal of queens could limit this potential as opposed to selecting towards increased queen longevity.

Immune priming and immune enhancement form the insect’s equivalent of the adaptive immune system found in vertebrates [115]. Both responses are defined as the enhanced protection resulting from past exposures to a pathogen [116] and incorporate memory of previous infections to either increase its efficacy in following exposures or enhance the antiviral response. Immune priming refers to responses specific to the pathogen whereas immune enhancement refers to responses non-specific to the pathogen [117]. Trans-generational immune priming has been described in honey bees for American foulbrood [118,119] along with its associated cost [120]. The reduced vertical transmission with age and between seasons for DWV and not for BQCV suggests that immune priming also occurs in honey bee queens within generations, as has been found in other insects [reviewed by 104,106,108]. A possible explanation could be the presence of RNA virus sequences that are produced during infections and serve as sources of siRNAs, even after clearance of the infection [111]. If so, this would imply that the diversity, with which honey bees counter viral stressors, is larger than previously thought. Possibly, the increased investment in immune priming could also explain the similar viral loads with increased infection frequencies in NSC. Comparing the increased virus resistance in honey bee queens [62,112,123] with tolerance in worker bees [39,45,46,108], and the influences both have on transmission pathways, emphasizes the importance of environmental and coevolutionary conditions on trait costs and trade-offs [54,55,124,125].

By focusing on the role of honey bee queens, this research adds to the growing literature on the relationship between viral infections and honey bee health. Evolutionary patterns of resistance and tolerance can form the theoretical foundation to incorporate virus resilience in breeding programs. A promising perspective as shown by the variability of vertical transmission over time, across queen ages and under different evolutionary conditions.

## Acknowledgements

We would like to thank Maria Bolt, Victor Kohut and Olivier Raujol for their participation in this study, as well as Viktor Deleu for her assistance in the lab. The SLU group would like to thank Åke Lyberg and Meghan Millbrath for help with the management of the colonies and the collection of the samples.

